# Three new genome assemblies of blue mussel lineages: North and South European *Mytilus edulis* and Mediterranean *Mytilus galloprovincialis*

**DOI:** 10.1101/2022.09.02.506387

**Authors:** Alexis Simon

**Affiliations:** ISEM, EPHE, IRD, Université Montpellier, Montpellier, France; Center of Population Biology and Department of Evolution and Ecology, University of California Davis, Davis, California, USA

**Keywords:** *Mytilus edulis*, *Mytilus galloprovincialis*, Genome assembly, 10X chromium

## Abstract

The blue mussel species complex (*Mytilus edulis*) is of particular interest both as model species in population genetics and ecology, but also as an economic resource in many regions. Using 10X genomics pseudo-long reads, I assembled genomes of three closely related blue mussel lineages from the *Mytilus* species complex in the Northern hemisphere. Given the huge diversity within and between lineages in this complex, the objective was to produce affordable genomic resources for population and evolutionary genomic studies to broaden the coverage of this diverse species complex. I used transcriptome guided corrections and scaffolding on a chromosome scale genome of a close species to reduce the fragmentation of the genomes. The result is a set of partially fragmented genomes of equivalent completeness to already published genomes. Three new draft genomes are added to the fast increasing genomic resources of this complex for the Mediterranean *M. galloprovlnclalls*, the South-European *M. edulis* and the the North-European *M. edulls*.

## 1 Rationale and objectives

The *Mytilus* species complex has been a model system in population genetics, adaptation, hybridization and speciation since genetic variants could be identified (Ahmad et al., 1977; Bierne et al., 2003; Fraïsse et al., 2016; Koehn & Mitton, 1972; Milkman & Beaty, 1970; Quesada, Wenne, et al., 1995; Simon et al., 2021; Skibinski et al., 1978). This genus is also an important food source worldwide. While *Mytilus* species are actively cultured, human selection has been limited in blue mussels as most of the production rely on naturally captured spat. Therefore, given the extreme diversity observed within and between species in this genus, extensive genomic resources are needed to both advance our fundamental knowledge and potentially accrue our capacity to improve its culture in the face of large environmental changes.

The *Mytilus* species complex is composed of three taxonomically recognized and partially reproductively isolated species in the northern hemisphere, *Mytilus edulis, M. galloprovincialis* and *M. trossulus;* and several other species in the southern hemisphere (*M. chilensis, M. platensis, M. planulatus*). Between those species, hybridization is possible in contact zones, either natural or anthropogenic. Within each species, evolutionary relevant lineages can be identified: two lineages in *M. galloprovincialis*, (i) an Atlantic lineage (MgalATL) and (ii) a Mediterranean lineage (MgalMED), separated by a hybrid zone at the Almeria-Oran front and Algerian coast (El Ayari et al., 2019; Fraïsse et al., 2016; Quesada, Zapata, et al., 1995), and three lineages in *M. edulis*, (i) an American lineage (MeduAM), (ii) a Southern European lineage (MeduEUS), and (iii) a Northern European lineage (MeduEUN) (Fraïsse et al., 2016; Simon et al., 2020). Two close species in the genus are of interest: *M. coruscus*, a species found in Asia, and *M. californianus*, a species found on the Pacific coast of North America. While they do not appear to readily hybridize with species of the *Mytilus* species complex, they are of similar interest to the study of marine mussels.

Increased effort has been put in the last few years to produce genomic resources in the *Mytilus* species complex. Quality genomes are available for the following lineages of *Mytilus:* Atlantic *M. galloprovincialis* (Gerdol et al., 2020), Southern European *M. edulis* (Corrochano-Fraile et al., 2022), *M. coruscus* (R. Li et al., 2020; Yang et al., 2021), *M. californianus* (Paggeot et al., 2022), and an American *M. edulis* assembly (unpublished, accession GCA_019925275). This increased effort has highlighted characteristics of interest in those species such as a high proportion of gene presence-absence (Gerdol et al., 2020) or evidence of genome duplication (Corrochano-Fraile et al., 2022).

As an effort to diversify the genomic resources available for the *Mytilus* species complex, I assembled and annotated the genomes of three lineages using the 10X chromium technology. While these assemblies were initially fragmented due to the high level of heterozygosity, I leveraged the existence of a chromosome scale assembly of a sister species to scaffold against it. I obtained assemblies equivalent to published ones in term of completeness for three new lineages of the Mytilus species complex using a low sequencing budget and publicly available data. The resources produced and the assembly pipeline are freely available for use by the community.

## 2 Methods

### General notes

The entire assembly was carried out using a Snakemake (Mölder et al., 2021) pipeline available on github at https://github.com/alxsimon/assembly_10x. Where deemed important, parameters are given in the text. For brevity and simplicity, not all information might be available in the text. However, all parameters, software versions and steps are retrievable from the repository.

### Important caveat

The assembled genome of MeduEUN *(M. edulis* Northern lineage) was initially thought to be *M. trossulus*. Therefore some assembly and annotation steps wrongly used *M. trossulus* transcriptomes. While this is not ideal, I think results have not been strongly impacted by this issue.

### 2.1 Biological material and DNA extraction

One individual for each species of interest was collected and processed fresh.

#### Collection locations

- *M. galloprovincialis* Mediterranean (MgalMED); Sete (43*°*24′35.3″*N*, 3*°*41′41.1″*E*, Occitanie, France, Mediterranean Sea)
- *M. edulis* Southern European (MeduEUS); Agon-Coutainville (49*°*0′44.784″*N*, 1*°*35′55.643″*W*, Normandy, France, English Channel)
- *M. edulis* Northern Europe (MeduEUN); Mishukovo (69*°*02′34.3″*N*, 33*°*01′45.9″*E*, Kola Bay, Russia, Barents Sea)

Whole mussels were placed in 50 mL falcon tubes containing 25 ml of TNES-Urea solution and incubated for 4-6 weeks at room temperature (TNES-Urea: 10 mM Tris-HCl pH 7.4; 120 mM NaCl; 10 mM EDTA pH 8.0; 0.5% SDS; 4 M urea).

After this period of pre-treatment at ambient temperature, proteinase K was added at a final concentration of 150 *^*g/mL and the solution was incubated overnight at 56*°*C.

High Molecular weight genomic DNA was extracted following Nakayama et al. (1994). A 15 mL Phase Lock Gel Heavy extraction was used with three steps of extraction using phenol/chloroform/isoamyl alcohol (25/24/1 proportions) equilibrated with Tris-HCl to pH 8.1, followed by two chloroform extractions. After the last extraction, the aqueous supernatant was precipitated with 2 volumes of 100% EtOH and the pellet was hooked from the solution with a sterile glass Pasteur pipette. The pellet was rinsed several times with 80% EtOH before being dried at room temperature.

DNA was resuspended with an appropriate volume of biomolecular water at 65rC for several hours, the duration of this incubation depending on the size of the granules. DNA was then stored at 4rC.

Prior to the construction of the DNA libraries, DNA was repaired with NEBNext FFPE DNA Repair Mix according to the manufacturer’s instructions.

### 2.2 Library preparation and sequencing

The 10X chromium library preparation and sequencing was subcontracted to the MGX platform (Montpellier, France). The 10X linked reads libraries for each individual were produced following the 10X Genomics Genome Reagen Kit *(Genome Solution)* protocol using a Chromium microfluidic chip. Libraries were subsequently sequenced on an Illumina NovaSeq 6000 with an S4 flowcell.

### 2.3 Preprocessing 10X reads

I preprocessed 10X reads using the following pipeline for use in several algorithms (section 2.4). I first removed duplicate reads using `Nubeam-dedup` (Dai and Guan, 2020; commit 25dd385). I then used `proc10xG` (https://github.com/ucdavis-bioinformatics/proc10xG; commit 7afbfcf) to split the reads from their 10X barcodes for further processing. I used a custom filtering step designed to remove under- and over-represented barcodes and their associated reads from the data (figs. S1 to S3, `filter_barcodes.R` script, personal communication of P-A Gagnaire). Reads were additionally filtered with `fastp` (v0.20.1; Chen et al., 2018) with the main objective to remove poly-G tails created by the Illumina Novaseq sequencing technology. I obtained at this point reads equivalent to a short-read sequencing run, usable in some parts of the assembly and quality control pipeline. Additionally, to obtain reads compatible to 10X genomics tools, I filtered and reassembled reads with their 10X barcodes in `proc10xG` using the `filter_10xReads.py` and `regen_10xReads.py` scripts.

### 2.4 Initial genome assemblies

For each genome I used `Supernova` v2.1.1 (Weisenfeld et al., 2017) to assemble raw 10X reads. To avoid hard stops in `Supernova` due to both data quantity slightly under 10X genomics recommendations and an overestimation of genome size by Supernova, I used all available reads (—`maxreads=`‘`all`’)and accepted extreme coverage (—`accept-extreme-coverage`). I produced every style of Fasta output available in `Supernova` but only used the pseudo-haploid output in the following pipeline.

To remove duplicate haplotypes in the assemblies, I followed the `purge_dups` pipeline (Guan et al., 2020; commit e1934bb). I first used the `longranger` v2.2.2 align algorithm to map preprocessed reads (section 2.3) to the pseudo-haploid genomes to use in the `ngscstat` step. `Minimap2` (v2.17; H. Li, 2017) was used in the contig self-mapping step. I obtained the purged assemblies using the `get_seqs` steps without restricting the purging to the end of contigs (without the `-e` option).

I used `AGOUTI` (https://github.com/svm-zhang/AGOUTI; commit a7e65d6; Zhang et al., 2016) to improve scaffolding by using paired end RNA-seq reads. For each species, a different set of published transcriptomes were used (see Supplementary File 1 for accession numbers). `AGOUTI` require a gene prediction as input in addition to RNA-seq reads. I used `Augustus` (v3.3.3; Stanke et al., 2008) to produce an intermediate annotation for each assembly using the *Caenorhabditis* model. RNA-seq reads were first cleaned using `rcorrector` (v1.0.4; Song and Florea, 2015) and `trimgalore` (v0.6.6; https://www.bioinformatics.babraham.ac.uk/projects/trim_galore; —`quality 20` —`stringency 1 -e 0.1` —`length 70`). RNA-seq reads were mapped independently using `bwa-mem2` (v2.2.1; Vasimuddin Md et al., 2019), and then merged as a common bam file for each assembly with `samtools` (v1.12; H. Li et al., 2009). Finally, the `AGOUTI` scaffolding pipeline was run using the gene prediction and mapped RNA-seq reads (`-minMQ 20 -maxFracMM 0.05`; using `python` v2.7.15 and `samtools` v1.10 for compatibility).

As a last step, I ran `Blobtoolkit` (v2.4.0; Challis et al., 2020) on the three assemblies to evaluate quality and potential contamination levels. I used a custom script (`btk_conta_extraction.py`) to filter the assembly contigs based on the taxids found by the `Blobtoolkit` pipeline to remove the most obvious contaminations. Contigs matching taxids associated with viruses, bacteria and non-mollusca eukaryotes were removed. More specifically contigs were removed for virus contamination if they presented a hit percentage of more than 10 % of their length. Contigs were removed for eukaryote contamination only when presenting only hits to taxa outside Mollusca on more than 10 % of their length. The list of retained contigs was used to filter the fasta assembly file using `seqkit` (v0.13.2; Shen et al., 2016).

### 2.5 Scaffolding on the *Mytilus coruscus* genome

At the time of assembly, the closest relative of the *Mytilus* species of interest with a chromosome scale assembly was *Mytilus coruscus* (GCA 017311375.1; Yang et al., 2021). Given the conserved number of 14 chromosomes in the *Mytilus* genus, I decided to scaffold our contigs on this high quality reference. Additionally, I had Oxford Nanopore reads for the MeduEUN individual. I used `minimap2` (v2.17; H. Li, 2017) and `LRScaf` (v1.1.10; https://github.com/shingocat/lrscaf) to first improve the MeduEUN assembly with this small amount of long reads.

Then for each assembly, I ran the scaffolder `RagTag` (v1.1.1; Alonge et al., 2021) to position contigs on the *M. coruscus* chromosomal assembly.

A final polishing step was performed using `Pilon` (v1.24; Walker et al., 2014). `Pilon` attempts to improve the assembly based on mapped read information (gap filling and error corrections). I used `bwa-mem2` to map the debarcoded and filtered 10X reads (section 2.3). In addition, Oxford Nanopore reads for MeduEUN were also used in `Pilon` for the given assembly.

### 2.6 Repeats

I masked repeats for the purpose of annotation using `RepeatModeler` (v2.0.1; Flynn et al., 2020) and `RepeatMasker` (v4.1.2-p1; Smit et al., 2013-2015) through the TETools DFAM container (v1.3.1; https://hub.docker.com/rZdfam/tetools). I first built repeat databases for each of five assemblies MgalMED, MeduEUS, MeduEUN, *M. coruscus* (GCA 017311375.1; Yang et al., 2021) and *M. galloprovincialis* from the Atlantic (GCA_900618805.1; Gerdol et al., 2020). Then, I built a common *Mytilus* database of repeats by merging those five databases using `cd-hit` (v4.8.1; Fu et al., 2012). I used the same options as used by default in `RepeatModeler`: `-aS 0.8 -c 0.8 -g 1 -G 0 -A 80 -M 10000`. Finally, soft repeat masking was performed on the assemblies with `RepeatMasker` using the merged database.

### 2.7 Annotation

I used `Braker2` (v2.1.6; Brna et al., 2021) as to obtain structural annotations, using both a protein database and RNA-seq reads (preprocessed in section 2.4). To build the protein database, I used all *Mollusca* proteins (taxid 6447) from OrthoDB (v10.1; Kriventseva et al., 2019). To provide gene presence hints, I mapped all RNA-seq reads for each species using `HISAT2` (v2.2.1; Kim et al., 2019).

I used the `Mantis` pipeline (v1.5.5; Queirós et al., 2021) to obtain consensus functional annotations of genes based on multiple databases. Protein sequences for each assembly were built from the structural annotations using the `python` module `gff3tool` (v2). `Mantis` was run with default parameters and databases: kofam (Aramaki et al., 2020), NPFM (Lu et al., 2020), eggNOG (Huerta-Cepas et al., 2019), pfam (El-Gebali et al., 2019), and tcdb (Saier et al., 2021).

### 2.8 NCBI submission

The NCBI submission process identified a few errors that needed correction. A small number of adaptor sequences and duplicates were removed to comply with NCBI requirements (see the rule `ncbi_submission_changes.smk` in the pipeline for more details). Assemblies are available under the following accessions: JAKGDF000000000 for MgalMED, JAKGDG000000000 for MeduEUS, and JAKGDH000000000 for MeduEUN.

### 2.9 Quality assessments and comparisons

Preprocessed 10X reads (section 2.3) were used to first estimate estimate genome size and heterozygosities of the three individuals. I used the reference free k-mer based method `GenomeScope` (https://github.com/tbenavi1/genomescope2.0; commit 5034ed4; Ranallo-Benavidez et al., 2020) and the fork of the `KMC` k-mer counting program (https://github.com/tbenavi1/KMC; commit 1df71f6).

Assembly statistics were computed with the python module `assembly_stats` (v0.1.4; Trizna, 2020).

To assess the remaining level of duplication in the assemblies, I used the program `KAT` (v2.4.2; Mapleson et al., 2017) to compare k-mer spectra from reads (preprocessed 10X) and from the assembly.

Finally, gene completion analyses were carried out using `BUSCO` (v5.1.1; Manni et al., 2021). I used both a Metazoan (`metazoa_odb10.2021-02-24`) and Molluscan (`mollusca_odb10.2020-08-05`) database to assess and compare new and published assemblies. I compared our assemblies to the following published ones:

- *M. coruscus*, GCA_017311375.1, Yang et al. (2021);
- *M. galloprovincialis* from the Altantic lineage, GCA_900618805.1, Gerdol et al. (2020);
- *M. edulis* from the Southern European lineage, GCA_905397895.1, Corrochano-Fraile et al. (2022)
- *M. edulis* from the American lineage, GCA_019925275.1.

### 2.10 Phylogenetic species tree

I compiled protein sequences from published *Mytilus* genomes and transcriptomes, in addition to the current three genomes and annotations, to build a species tree. I used transcriptomes produced in Popovic and Riginos (2020) for American *M. edulis*, Mediterranean *M. galloprovincialis, M. trossulus, M. californianus*. Transcriptomes were translated using the `seqkit` program (v2.2.0; Shen et al., 2016) I used genomes and associated annotations of *M. coruscus* (GCA 017311375.1; Yang et al., 2021), Southern European *M. edulis* (GCA_905397895.1; Corrochano-Fraile et al., 2022), Atlantic *M. galloprovincialis* (GCA_900618805.1; Gerdol et al., 2020), and American *M. edulis* (GCA_019925275.1; annotation as personal communication of Tiago Hori, PEIMSO). For GCA_019925275.1, protein sequences where retrieve from the fasta and gff files using the python module gff3tool (v2.1.0).

I used `OrthoFinder` (v2.5.4; Emms and Kelly, 2015, 2019) to find orthogroups and orthologue genes. Species tree was inferred using the MSA method of `OrthoFinder` (Emms & Kelly, 2018) using the `MAFFT` aligner (v7.505; Katoh and Standley, 2013), the `STRIDE` species tree rooting algorithm (Emms & Kelly, 2017) and the `FastTree` software for tree inference (v2.1.11; Price et al., 2009).

## 3 Results and discussion

### 3.1 Assembly results

I introduce here newly assembled genomes from three lineages of the *Mytilus* species complex. With a limited sequencing budget of *∼* 3000 per genome, I managed to produce draft genomes of enough quality to be useful in applications such as population genomics and genetics. The method of 10X chromium pseudo-long reads combined with scaffolding using published data provided a quality comparable to published assemblies for *Mytilus* species.

The pre-assembly k-mer analysis carried out using GenomeScope (using 21-mers) showed that, as expected, the genomes of MgalMED and MeduEUN were highly heterozygous with values of 3.49 and 4.03 % respectively (fig. 1). The heterozygous peak of MeduEUS was not identifiable due to lower sequencing depth and GenomeScope provides a bad quality fit that I did not interpret.

**Figure 1:**
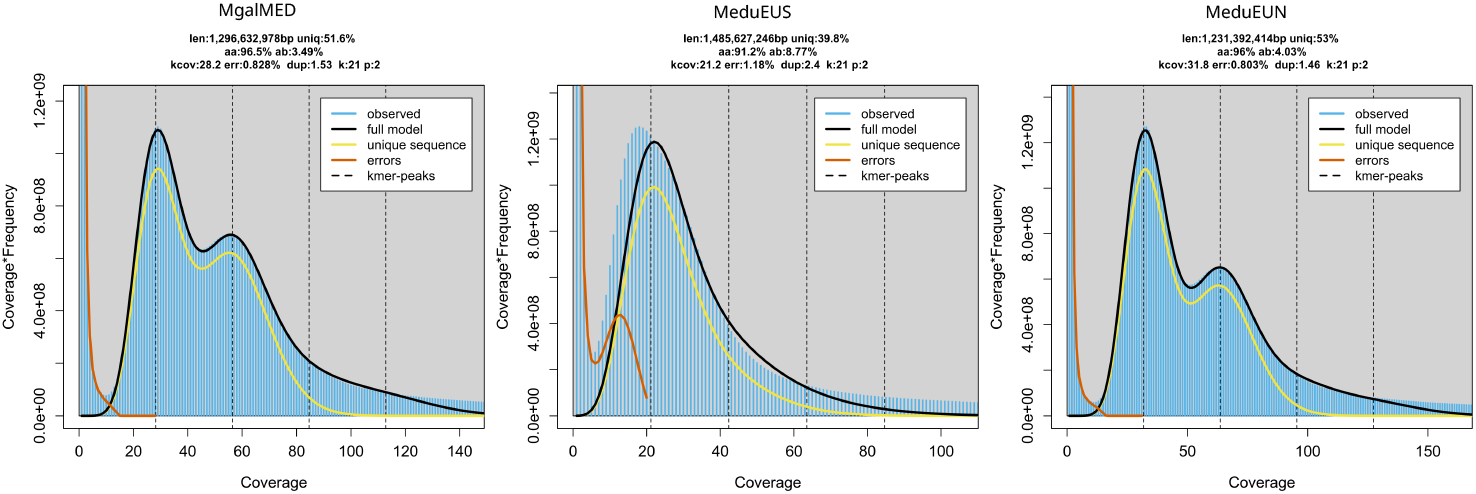
k-mer profile plots computed with 21-mers using GenomeScope. Coverage histogram of the k-mers in blue. Lines represent the fit of the GenomeScope models. len: inferred genome length; uniq: percentage of the genome that is unique, aa: overall homozigosity; ab: overall heterozygosity; kcov: mean k-mer coverage for heterozygous bases; err: reads error rate; dup: average rate of read duplication; k: k-mer size; p: ploidy.

Highly heterozygous genomes brings the risk of having a large number of duplicate contigs due to the separate assembly of the multiple alleles from given locus. For this reason, I used the `purge_dups` pipeline to try removing a maximum of such bias. This procedure reduced the number of complete duplicated genes in all assemblies (fig. S5, v1 to v4). Overall, the compared `KAT` spectra analyses show that the assembly steps reduced the amount of duplication in all assemblies (fig. S4).

Assembly statistics for the three new assemblies and four publicly available assemblies are presented in table 1 (complete statistics are presented in Supplementary File 2). With the exception of MeduEUS, GC content and genome sizes are close to what is found in other lineages. While the number of scaffolds for the new assemblies is still high, most of the genomes are contained in 14 large scaffolds corresponding to the number of chromosomes found in *Mytilus* species. Residual scaffolds are sequences that could not be placed on the *M. coruscus* genome, maybe due to inter-species presence-absence or too high divergence.

**Table 1:** Assembly statistics comparisons. C for contigs and S for scaffolds.

| assembly | C.L50 | C.N50 | C.median | C.sequence_count | S.L50 | S.N50 | S.gc_content | S.median | S.sequence_count | S.total_bps |
| --- | --- | --- | --- | --- | --- | --- | --- | --- | --- | --- |
| Mcor_GCA017311375 | $2.61 \times 10^2$ | $1.48 \times 10^6$ | $4.21 \times 10^4$ | 6449 | 6 | $9.95 \times 10^7$ | 32.4 | $2.13 \times 10^4$ | 4434 | $1.57 \times 10^9$ |
| MgalATL_GCA900618805 | $4.92 \times 10^3$ | $7.70 \times 10^4$ | $3.95 \times 10^4$ | 22922 | 1903 | $2.08 \times 10^5$ | 32.1 | $7.76 \times 10^4$ | 10577 | $1.28 \times 10^9$ |
| MeduEUS_GCA905397895 | $9.77 \times 10^2$ | $5.11 \times 10^5$ | $1.85 \times 10^5$ | 5966 | 464 | $1.10 \times 10^6$ | 32.2 | $2.92 \times 10^5$ | 3339 | $1.83 \times 10^9$ |
| MeduAM_GCA019925415 | $8.35 \times 10^2$ | $4.91 \times 10^5$ | $6.25 \times 10^4$ | 9686 | 6 | $1.17 \times 10^8$ | 32.3 | $2.89 \times 10^4$ | 1119 | $1.65 \times 10^9$ |
| MgalMED_v7 | $1.69 \times 10^4$ | $2.63 \times 10^4$ | $7.48 \times 10^3$ | 115913 | 9 | $7.20 \times 10^7$ | 32.1 | $4.53 \times 10^3$ | 41122 | $1.66 \times 10^9$ |
| MeduEUS_v7 | $2.45 \times 10^4$ | $2.36 \times 10^4$ | $6.48 \times 10^3$ | 169324 | 415 | $2.28 \times 10^5$ | 34.6 | $5.00 \times 10^3$ | 73616 | $2.08 \times 10^9$ |
| MeduEUN_v7 | $1.59 \times 10^4$ | $2.92 \times 10^4$ | $8.96 \times 10^3$ | 105892 | 9 | $7.71 \times 10^7$ | 32.1 | $6.16 \times 10^3$ | 28220 | $1.76 \times 10^9$ |

To assess the completeness of our assemblies, I compared them to four *Mytilus* published assemblies on the basis of a Metazoan and a Molluscan set of single copy orthologous genes using BUSCO (fig. 2). Overall, I show that our assemblies are equivalent to the published ones in terms of completeness.

**Figure 2:**
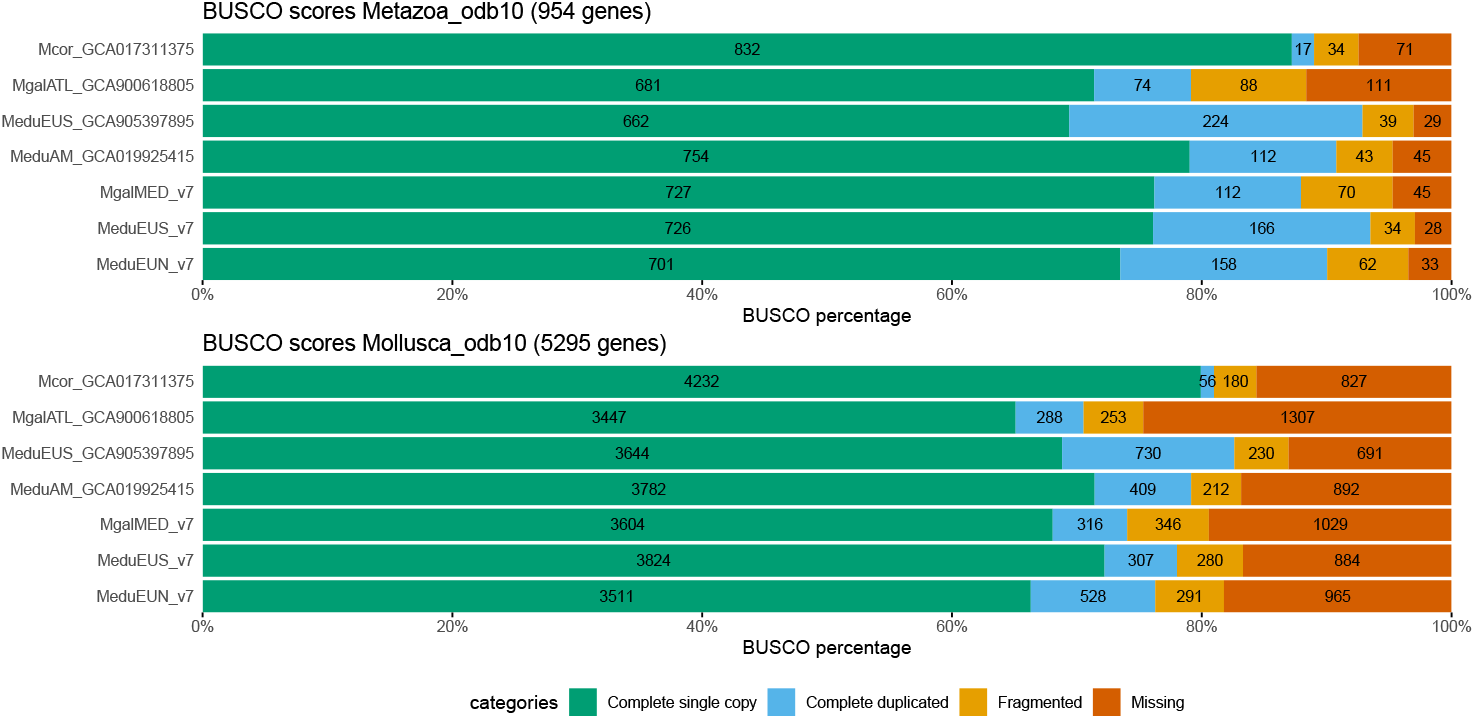
Busco scores

The MeduEUS assembly suffered from an increased level of contamination compared to the other two assemblies. Despite a broad contaminant filtering step, the Busco blob analysis (fig. S7) shows a second GC content peak centered around 40% (above the 32% peaks of this and other genomes). This contamination could for instance be due to bacteria present in the mussel if its physiological state was degraded before fixation. The metagenomic composition of contaminants was not investigated.

### 3.2 Repeat contents

More than half of each genome was identified as repeats and masked by `RepeatMasker`. Repeats are estimated to amount to 57.22, 61.10 and 55.53% of MgalMED, MeduEUS and MeduEUN assemblies respectively. The majority of repeats are unclassified, followed by retroelements with a balanced contribution of LINEs and LTR elements (detailed `RepeatMasker` results in table S2).

### 3.3 Annotations

The assemblies were annotated using a consensus pipeline to provide additional resources for future uses. These annotation were used to produce the following species tree.

### 3.4 Species tree

OrthoFinder assigned 499004 genes (91.9% of total) to 50543 orthogroups. Fifty percent of all genes were in orthogroups with 13 or more genes (G50 was 13) and were contained in the largest 11982 orthogroups (O50 was 11982). There were 1852 orthogroups with all species present and 173 of these consisted entirely of single-copy genes.

The species tree was built using 1300 orthogroups with a minimum of 81.8% of species having single-copy genes in any orthogroup (fig. 3). It shows that the assemblies and the published genomes and transcriptomes are clustering as expected.

**Figure 3:**
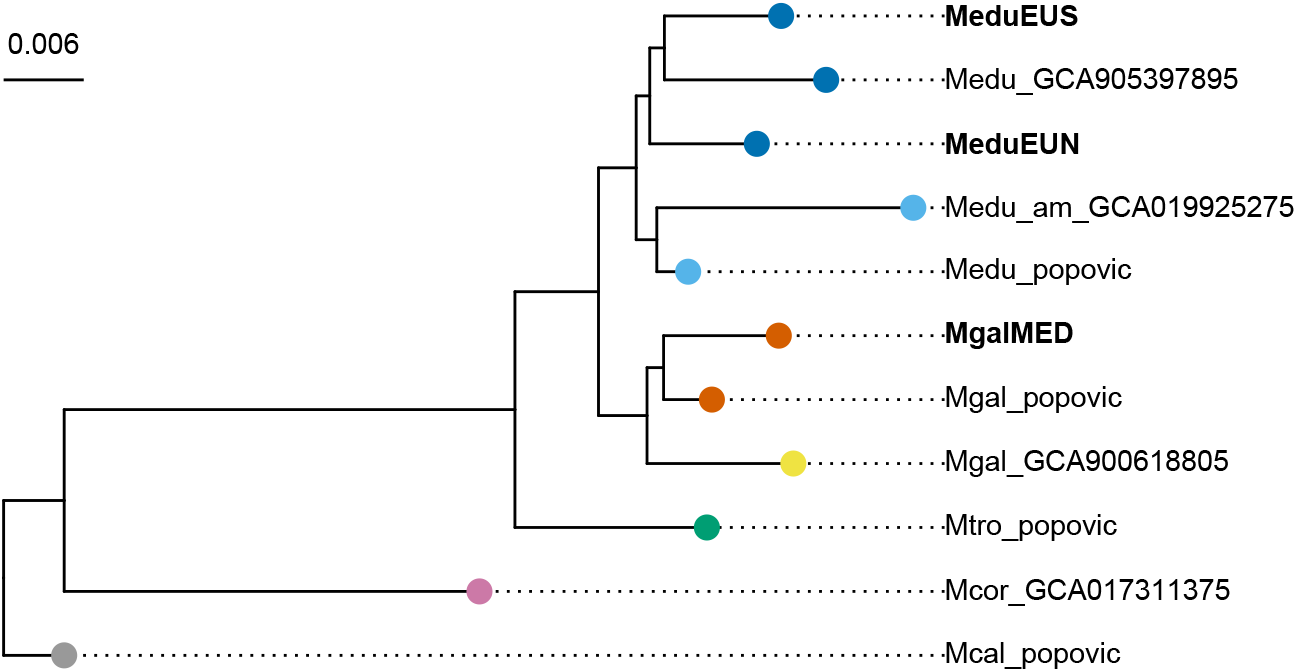
Species tree using 1300 orthogroups with a minimum of 81.8% of species having single-copy genes in any orthogroup. Color coding - dark blue: *M. edulis* Europe, light blue: *M. edulis* America, red: *M. galloprovincialis* Mediterranean Sea, yellow: *M. galloprovincialis* Atlantic, green: *M. trossulus*, pink: *M. coruscus*, gray: *M. californianus*. The “popovic” suffix indicate transcriptome date from Popovic and Riginos (2020).

## Supporting information

Supp_file_1_RNAseq_accessions.tsv

Supp_file_2_assembly_stats.tsv

## Data availability

Raw data are available under BioProject PRJNA785550. Assemblies are available under accessions JAKGDF000000000 (MgalMED), JAKGDG000000000 (MeduEUS), and JAKGDH000000000 (MeduEUN). The assembly pipeline is available at https://github.com/alxsimon/assembly_10x. Annotations, and the OrthoFinder pipeline and results are available in the Zenodo archive 10.5281/zenodo.7034399. Data and code for this manuscript are available at https://github.com/alxsimon/Mytilus_ref_genomes_paper.

## Acknowledgements

I thank Nicolas Bierne and Pierre-Alexandre Gagnaire for their help on the project and discussions; Christine Arbiol for DNA extractions; Maurine Hammel and Erika Burioli for sharing their transcriptome data; and Petr Strelkov for sampling of the *M. trossulus*. Some bioinformatic analyses were performed on the Core Cluster of the Institut Frangais de Bioinformatique (ANR-11-INBS-0013). I thank Tiago Horri for providing his annotation of the American *Mgtilus edulis* genome (GCA_019925275.1). AS was supported by the French ANR grant TRANSCAN (ANR-18-CE35-0009) and by the National Institute of General Medical Sciences of the National Institutes of Health (grants NIH R01 GM108779 and R35 GM136290 to Graham Coop).

## Supplementary Information

**Table S1:**
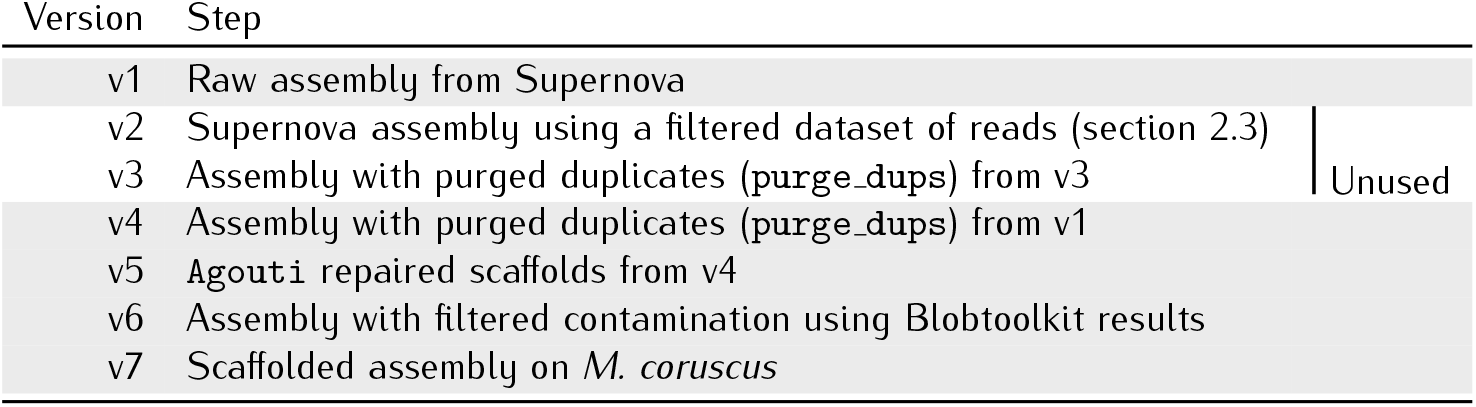
Steps carried out for each version of assembly

**Figure S1:**
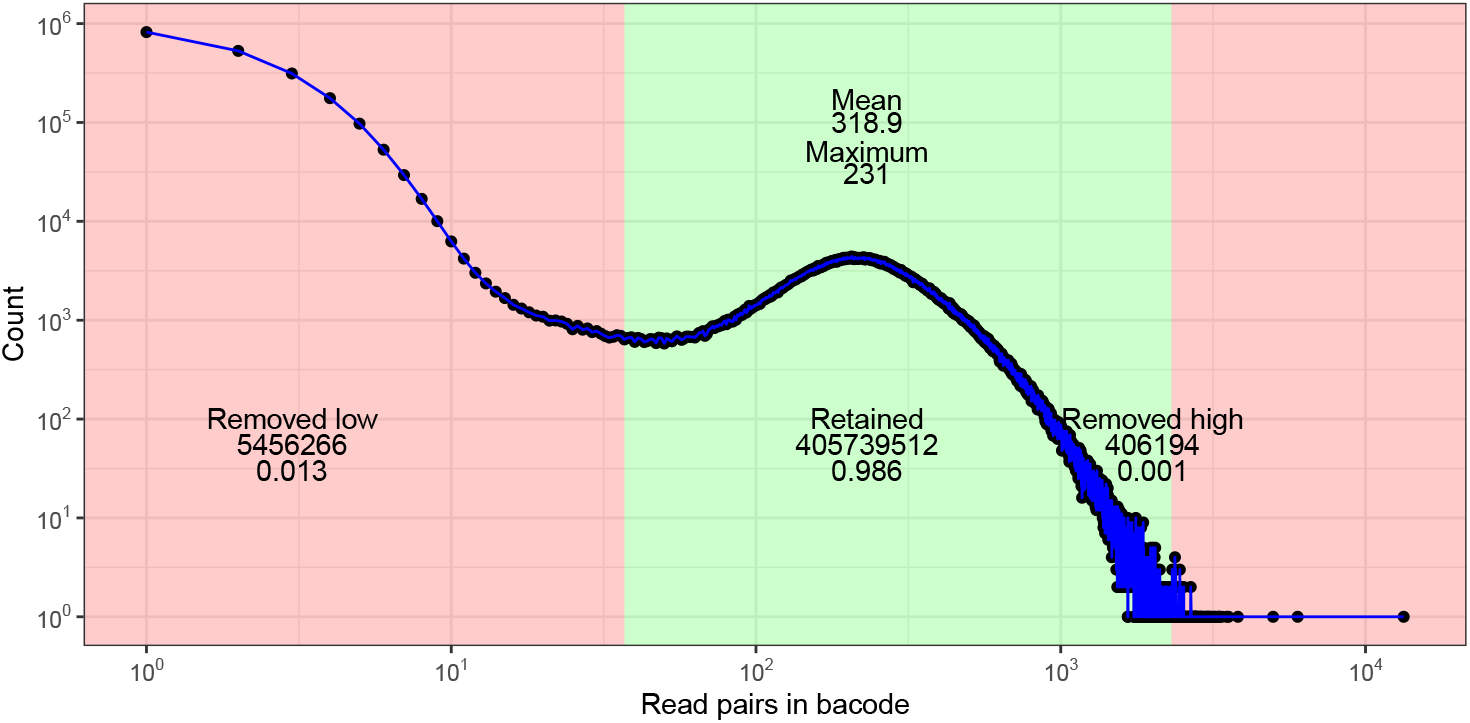
Histogram of read pairs identified for each 10X barcode for MgalMED. As part of preprocessing step, barcodes for which too few or too many read pairs are associated with each unique barcode are removed from the dataset (red regions).

**Figure S2:**
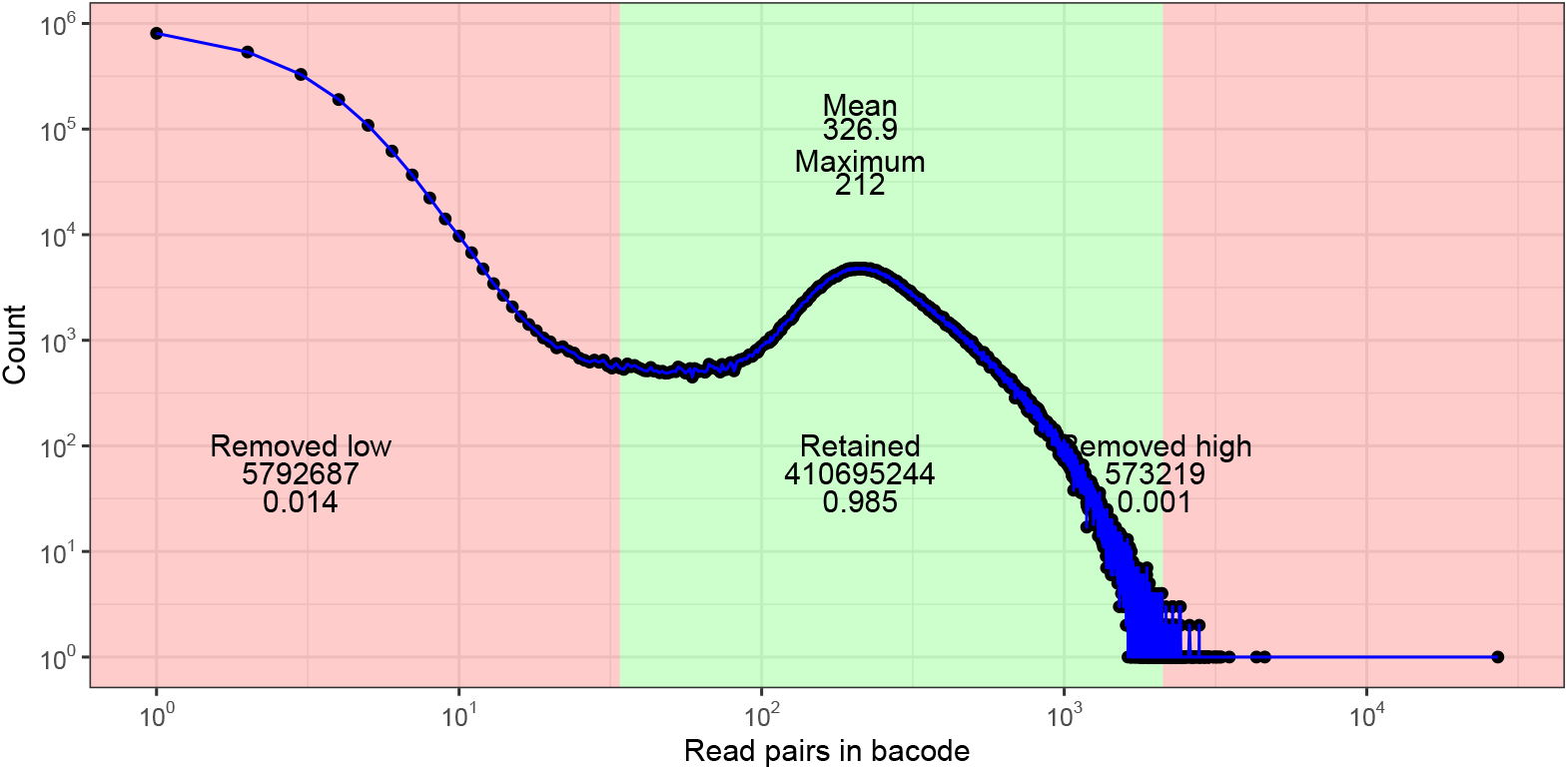
Histogram of read pairs identified for each 10X barcode for MeduEUS. As part of preprocessing step, barcodes for which too few or too many read pairs are associated with each unique barcode are removed from the dataset (red regions).

**Figure S3:**
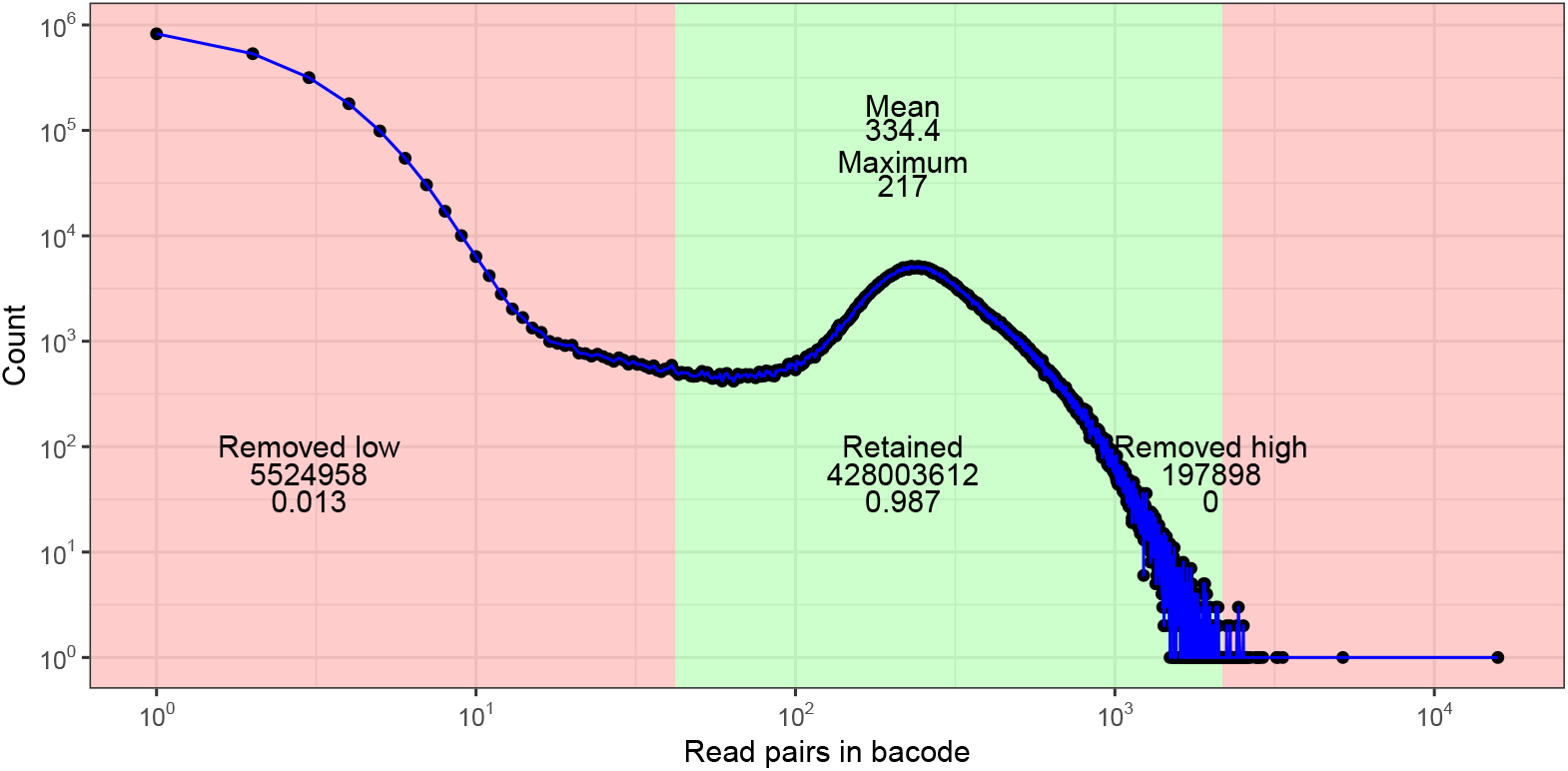
Histogram of read pairs identified for each 10X barcode for MeduEUN. As part of preprocessing step, barcodes for which too few or too many read pairs are associated with each unique barcode are removed from the dataset (red regions).

**Figure S4:**
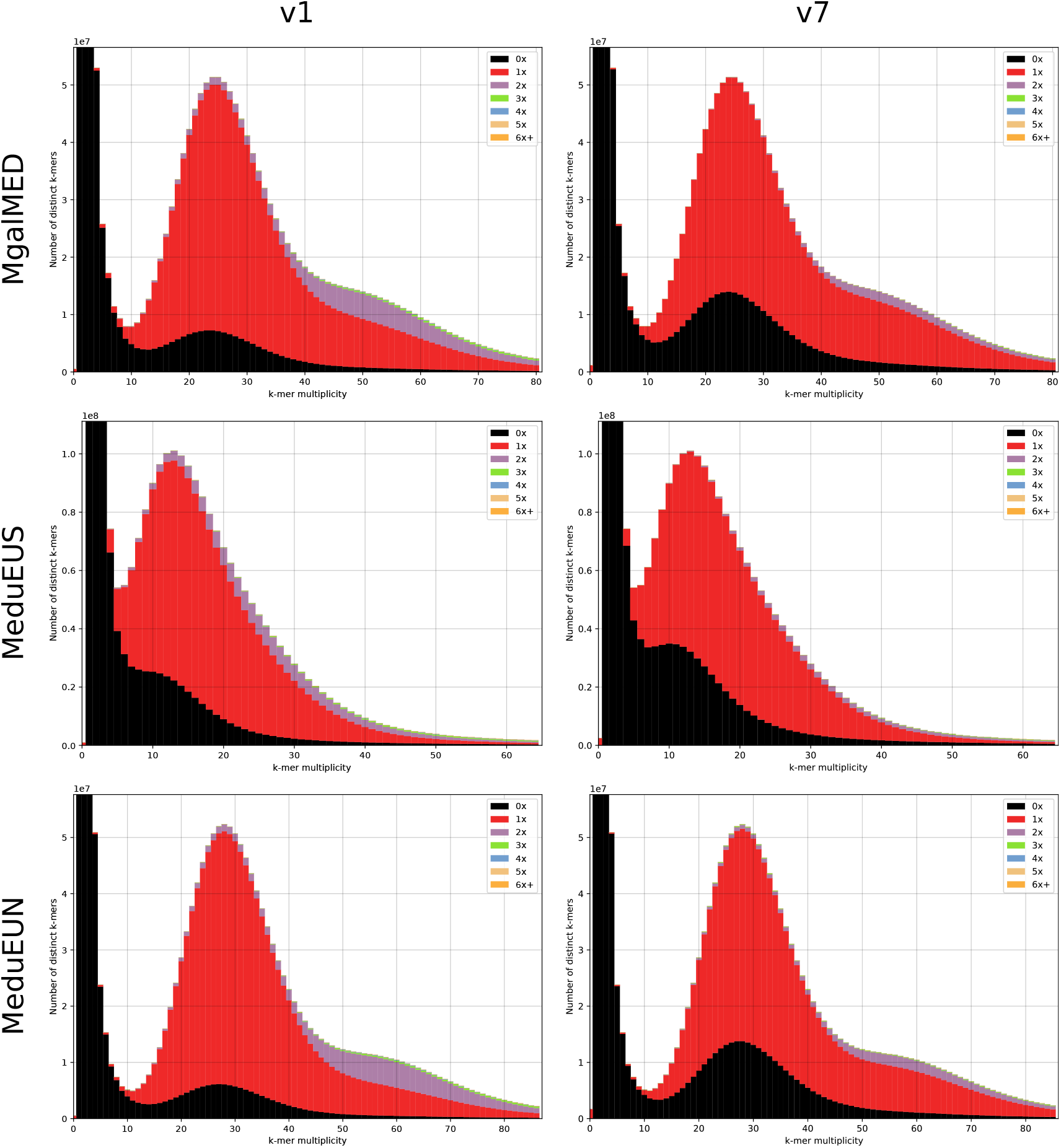
KAT comparison of k-mer spectra between assembly and preprocessed reads. Results are compared between the initial assembly (v1, left column) and the final assembly (v7, right column) for each sample MgalMED, MeduEUS and MeduEUN (from top to bottom).

**Figure S5:**
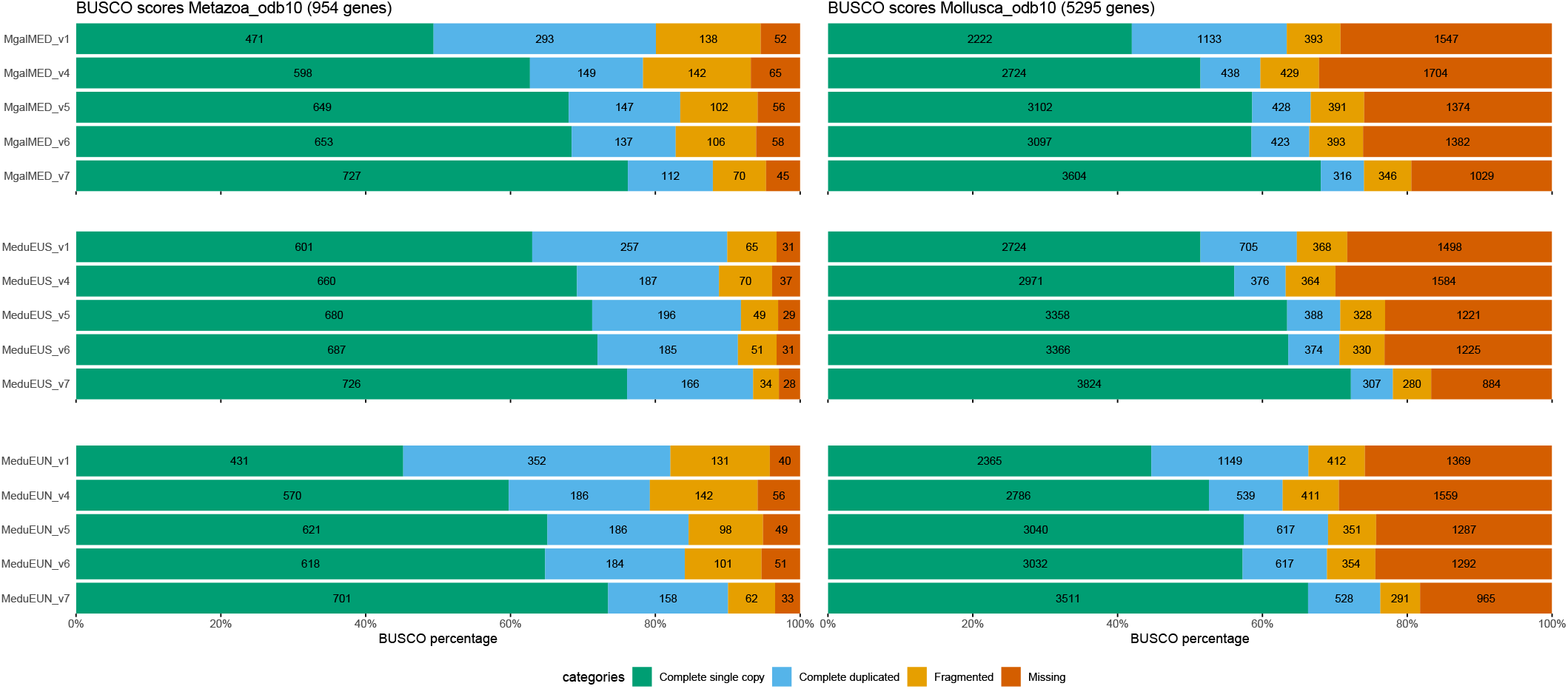
Busco scores on the metazoa_odb10 (left panels) and mollusca_odb10 (right panels) databases for each assembly MgalMED, MeduEUS and MeduEUN (top to bottom panels) across several assembly versions (v1, v4, v5, v6, v7).

**Figure S6:**
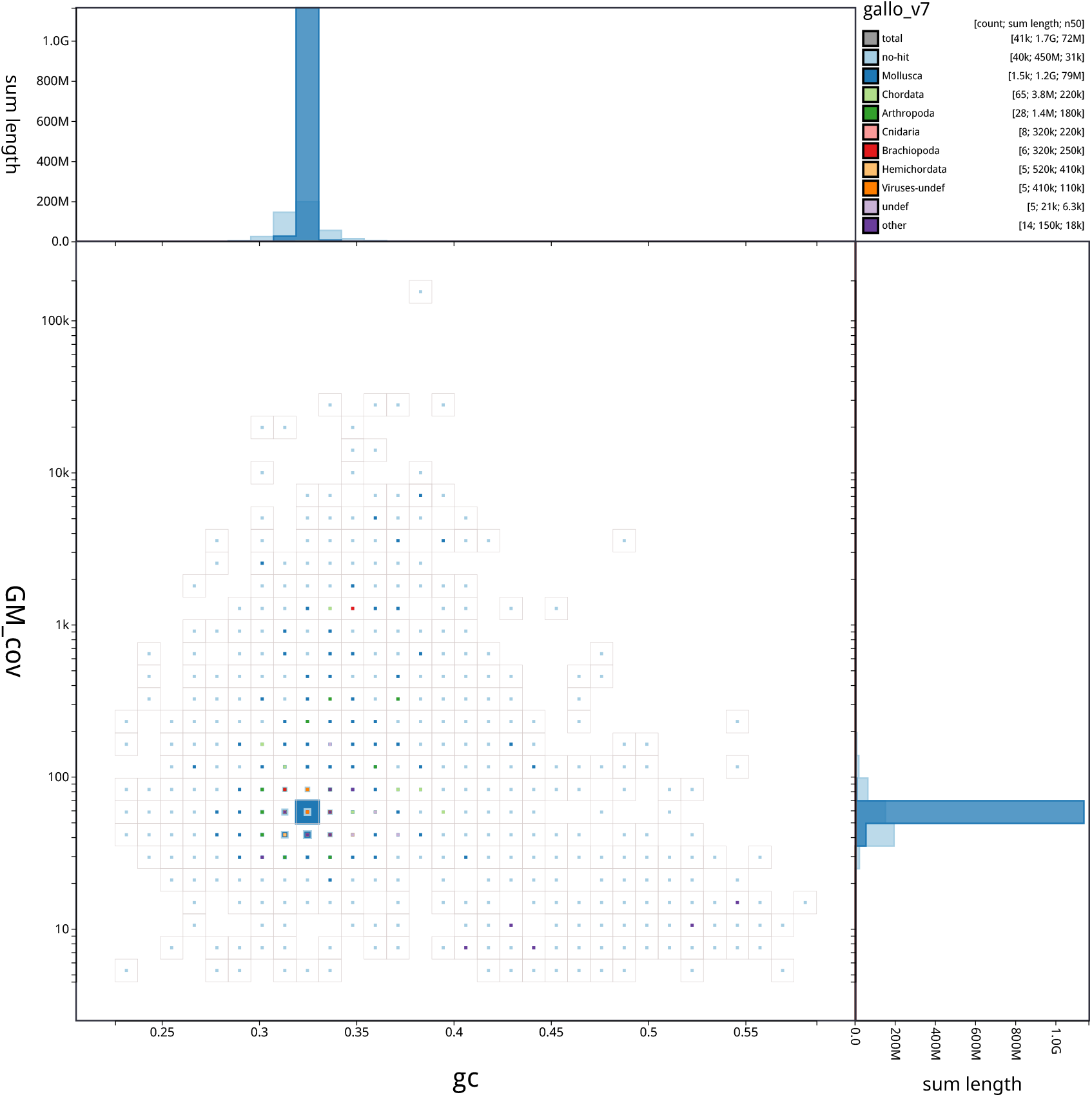
Blobtoolkit blob plot of base coverage in GM against GC proportion for scaffolds in assembly MgalMED_v7. Scaffolds are colored by phylum and binned at a resolution of 30 divisions on each axis. Colored squares within each bin are sized in proportion to the sum of individual scaffold lengths on a square-root scale, ranging from 1,005 to 1,127,105,406. Histograms show the distribution of scaffold length sum along each axis.

**Figure S7:**
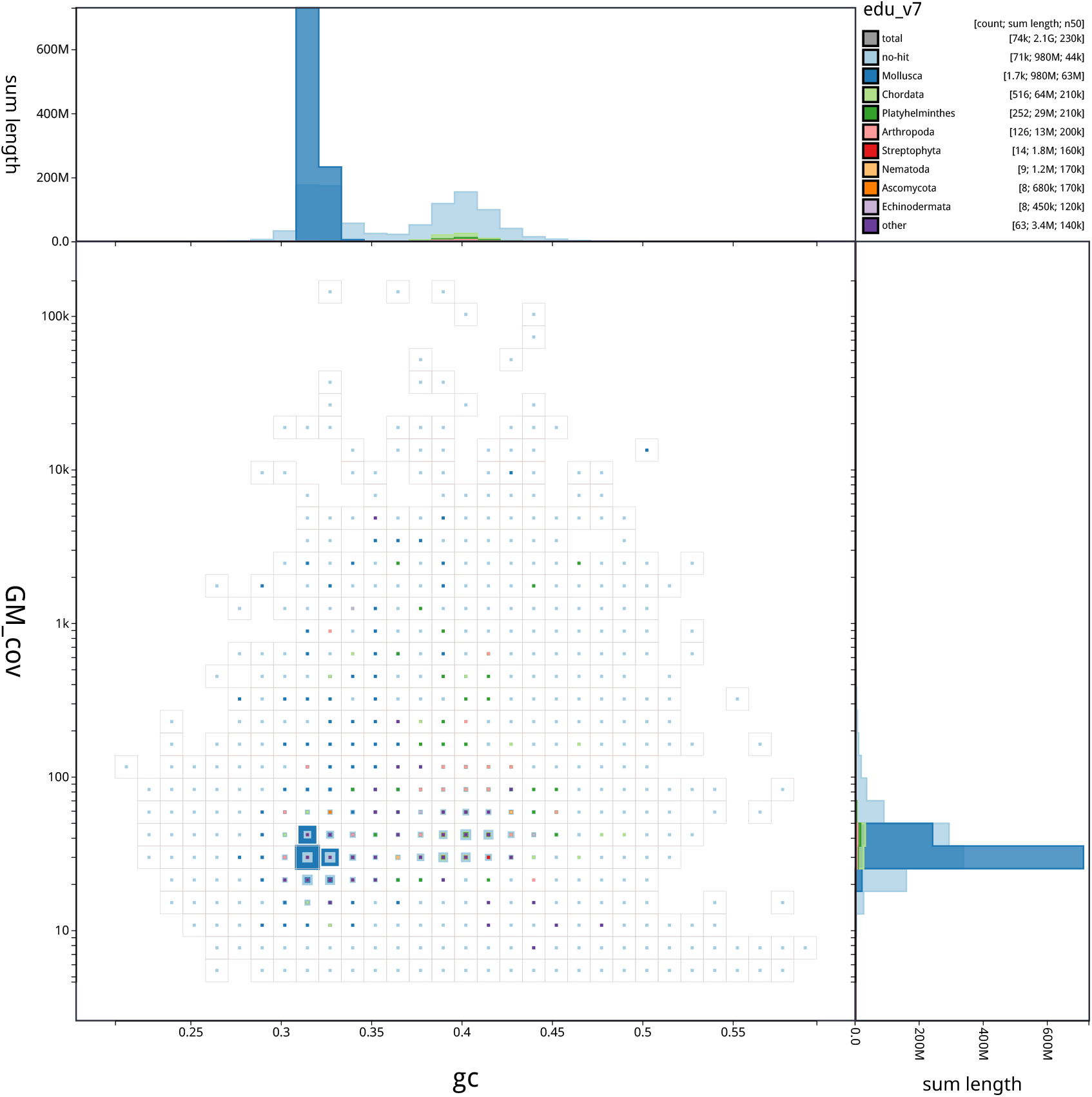
Blobtoolkit blob plot of base coverage in GM against GC proportion for scaffolds in assembly MeduEUS_v7. Scaffolds are colored by phylum and binned at a resolution of 30 divisions on each axis. Colored squares within each bin are sized in proportion to the sum of individual scaffold lengths on a square-root scale, ranging from 987 to 487,517,891. Histograms show the distribution of scaffold length sum along each axis.

**Figure S8:**
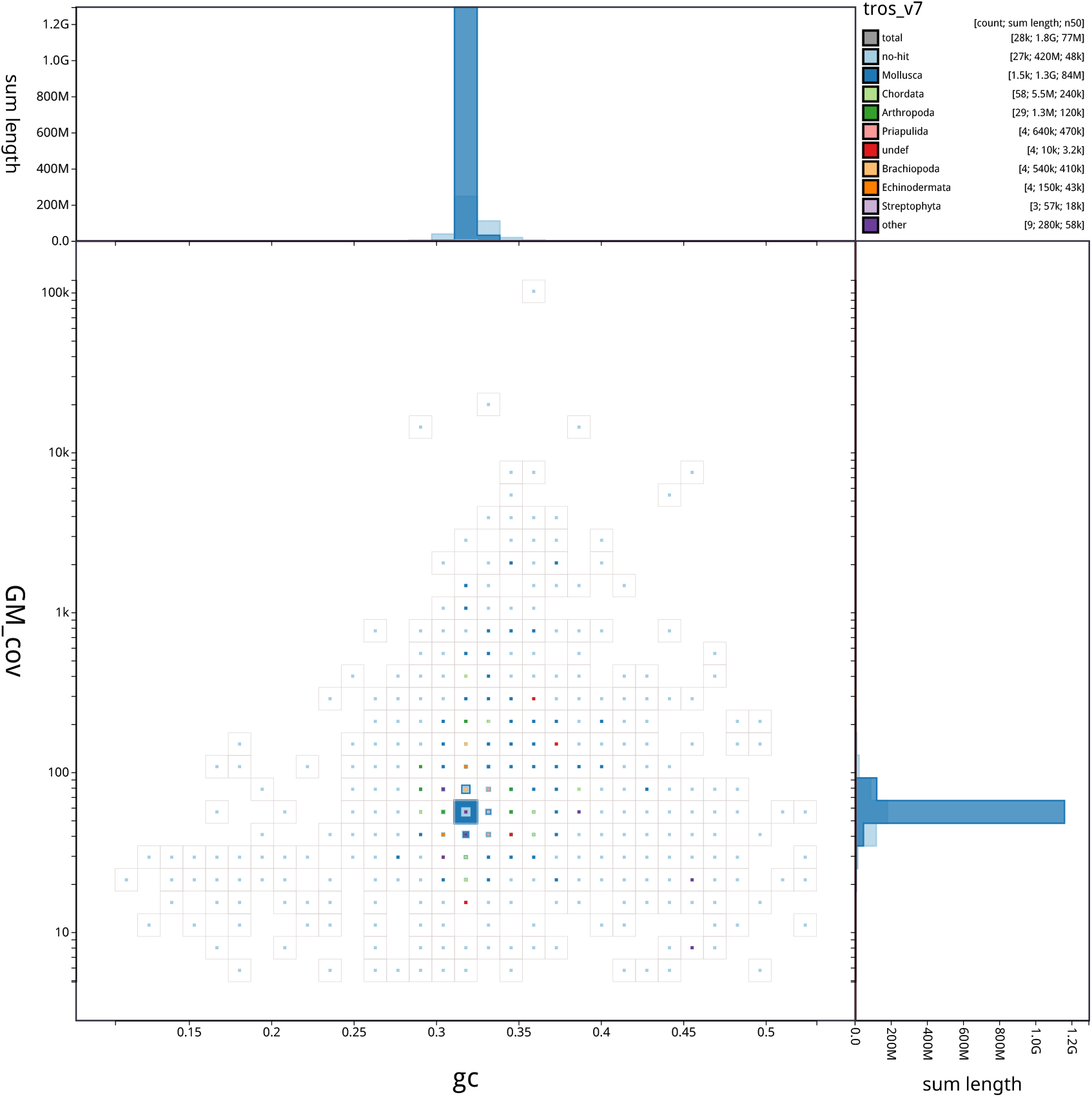
Blobtoolkit blob plot of base coverage in GM against GC proportion for scaffolds in assembly MeduEUN_v7. Scaffolds are colored by phylum and binned at a resolution of 30 divisions on each axis. Colored squares within each bin are sized in proportion to the sum of individual scaffold lengths on a square-root scale, ranging from 1,011 to 1,136,301,196. Histograms show the distribution of scaffold length sum along each axis.

**Figure S9:**
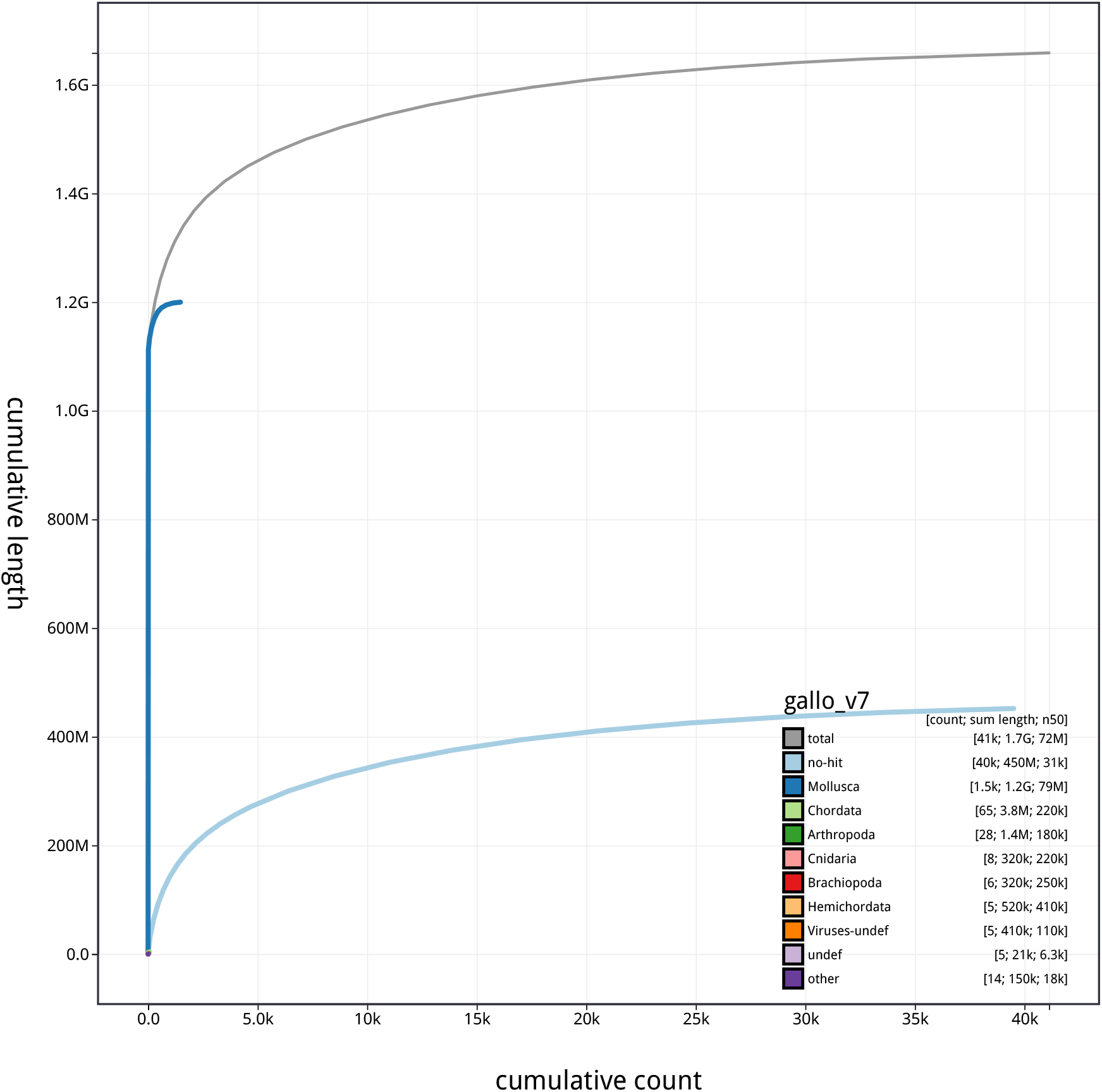
Blobtoolkit cumulative scaffold length for assembly MgalMED_v7. The gray line shows cumulative length for all scaffolds. Colored lines show cumulative lengths of scaffolds assigned to each phylum using the bestsumorder taxrule.

**Figure S10:**
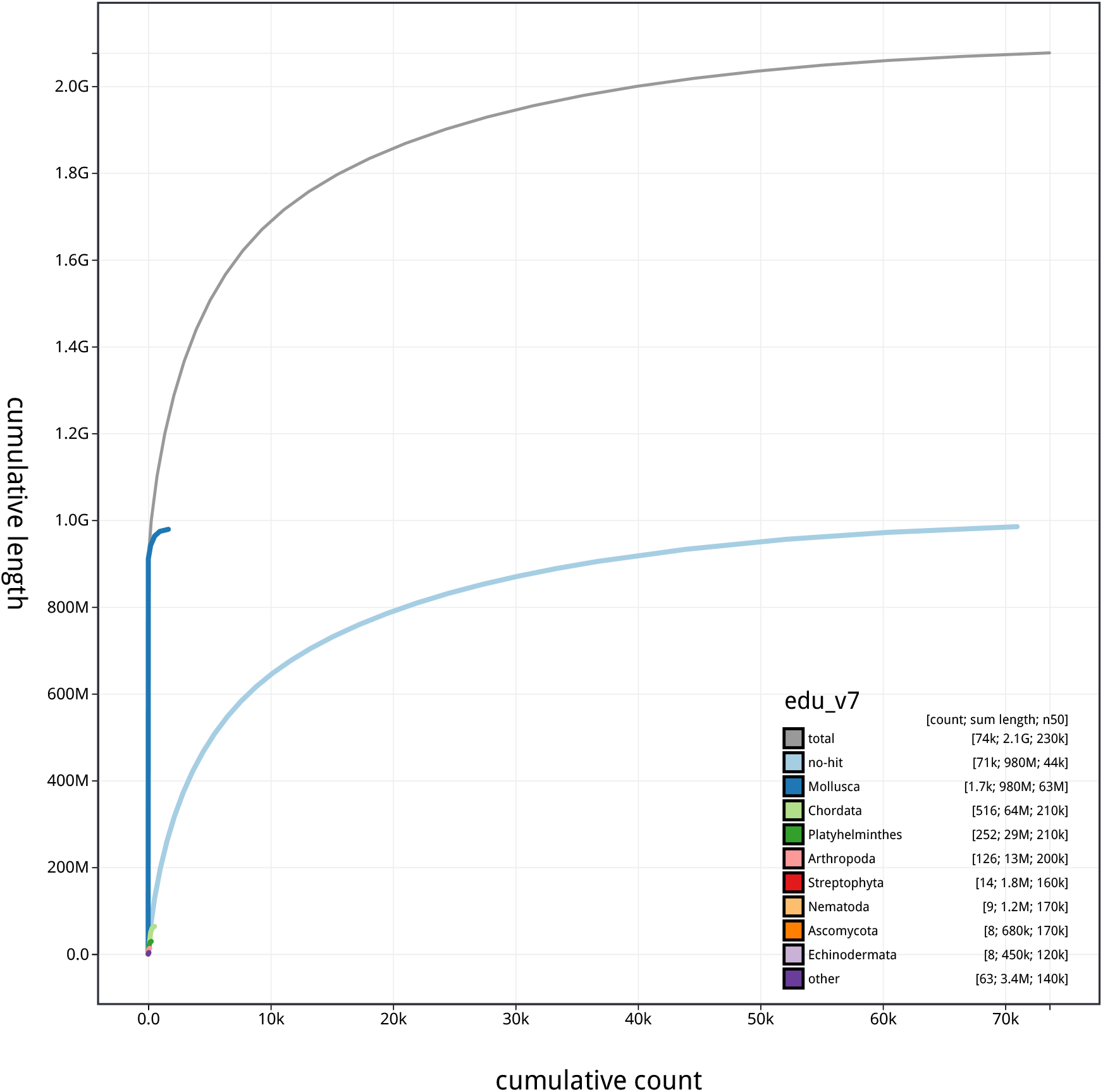
Blobtoolkit cumulative scaffold length for assembly MeduEUSv7. The gray line shows cumulative length for all scaffolds. Colored lines show cumulative lengths of scaffolds assigned to each phylum using the bestsumorder taxrule.

**Figure S11:**
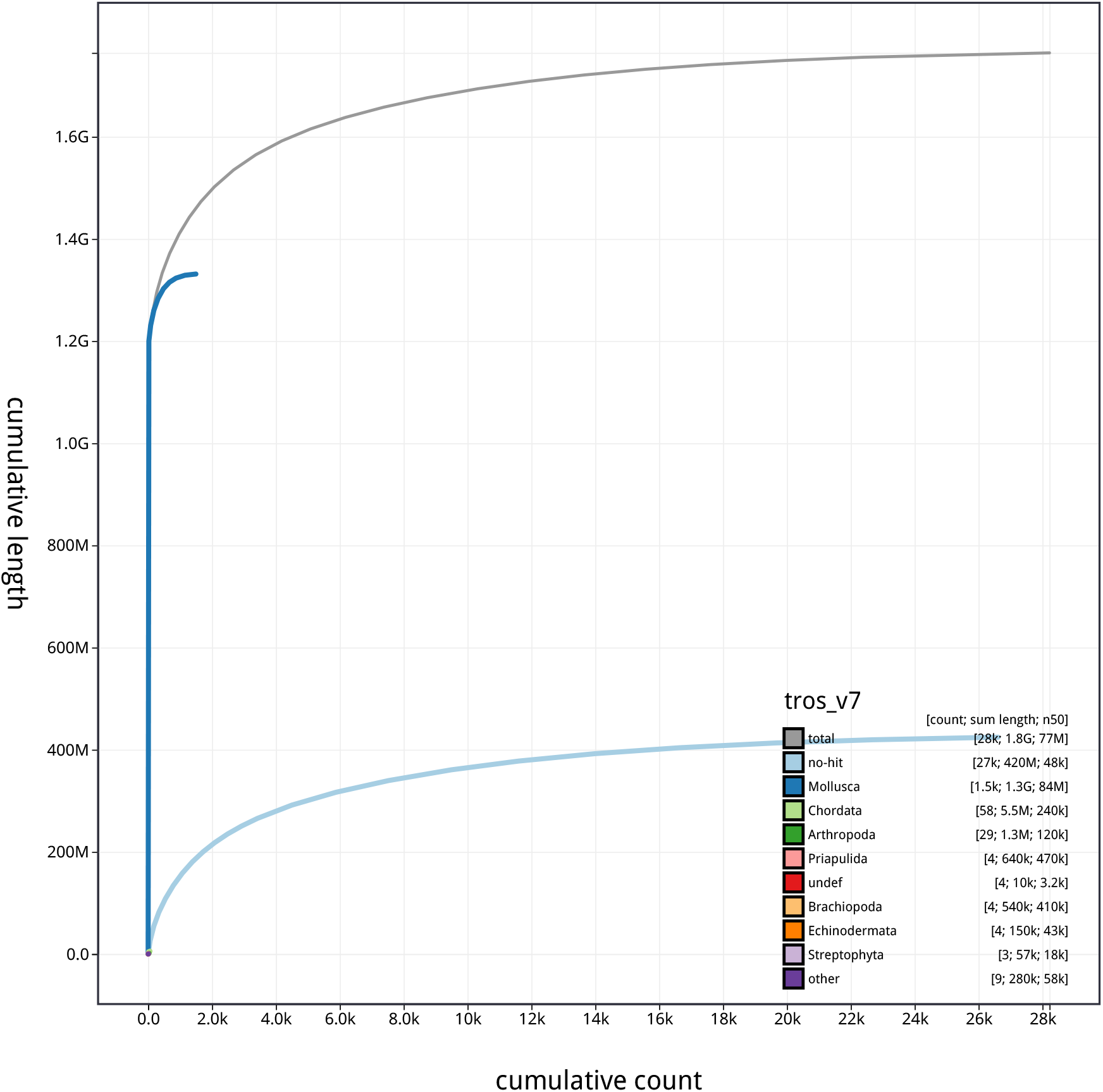
Blobtoolkit cumulative scaffold length for assembly MeduEUN_v7. The gray line shows cumulative length for all scaffolds. Colored lines show cumulative lengths of scaffolds assigned to each phylum using the bestsumorder taxrule.

**Figure S12:**
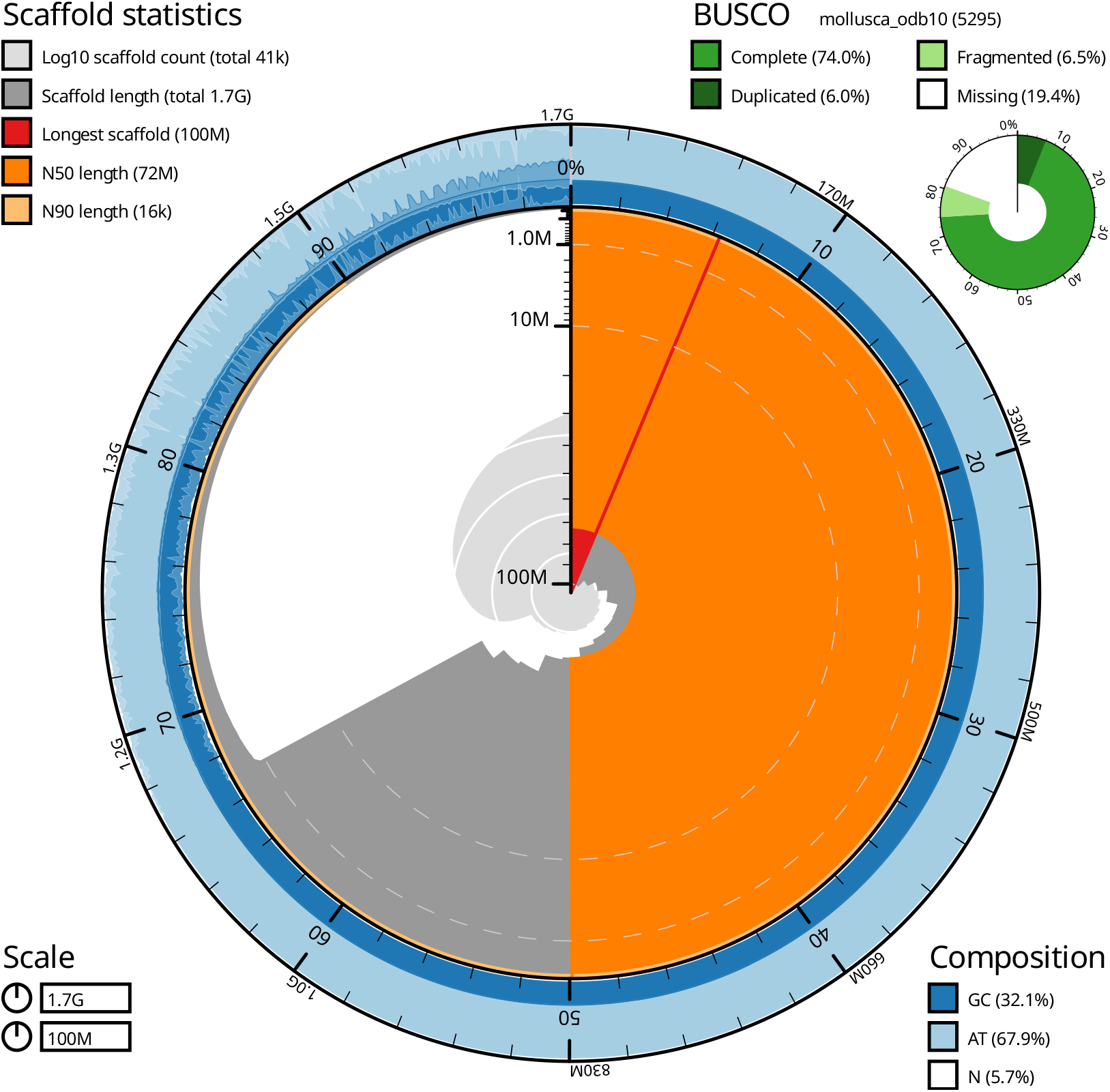
Blobtoolkit snail plot summary of assembly statistics for assembly MgalMED_v7. The main plot is divided into 1,000 size-ordered bins around the circumference with each bin representing 0.1% of the 1,658,656,017 bp assembly. The distribution of scaffold lengths is shown in dark gray with the plot radius scaled to the longest scaffold present in the assembly (104,729,878 bp, shown in red). Orange and pale-orange arcs show the N50 and N90 scaffold lengths (71,952,646 and 16,011 bp), respectively. The pale gray spiral shows the cumulative scaffold count on a log scale with white scale lines showing successive orders of magnitude. The blue and pale-blue area around the outside of the plot shows the distribution of GC, AT and N percentages in the same bins as the inner plot. A summary of complete, fragmented, duplicated and missing BUSCO genes in the mollusca_odb10 set is shown in the top right.

**Figure S13:**
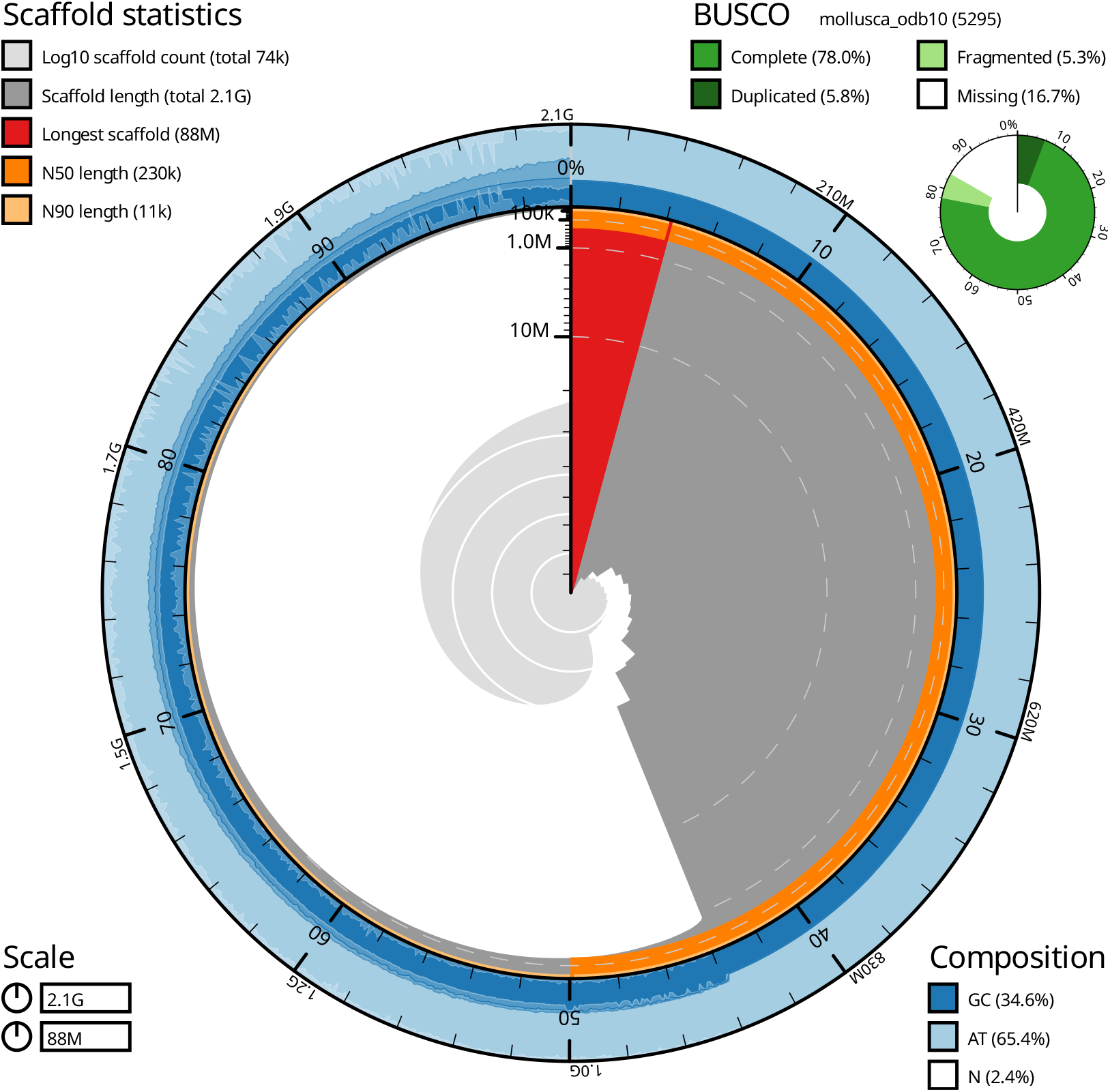
Blobtoolkit snail plot summary of assembly statistics for assembly MeduEUSv7. The main plot is divided into 1,000 size-ordered bins around the circumference with each bin representing 0.1% of the 2,076,685,641 bp assembly. The distribution of scaffold lengths is shown in dark gray with the plot radius scaled to the longest scaffold present in the assembly (88,305,666 bp, shown in red). Orange and pale-orange arcs show the N50 and N90 scaffold lengths (228,263 and 10,773 bp), respectively. The pale gray spiral shows the cumulative scaffold count on a log scale with white scale lines showing successive orders of magnitude. The blue and pale-blue area around the outside of the plot shows the distribution of GC, AT and N percentages in the same bins as the inner plot. A summary of complete, fragmented, duplicated and missing BUSCO genes in the mollusca_odb10 set is shown in the top right.

**Figure S14:**
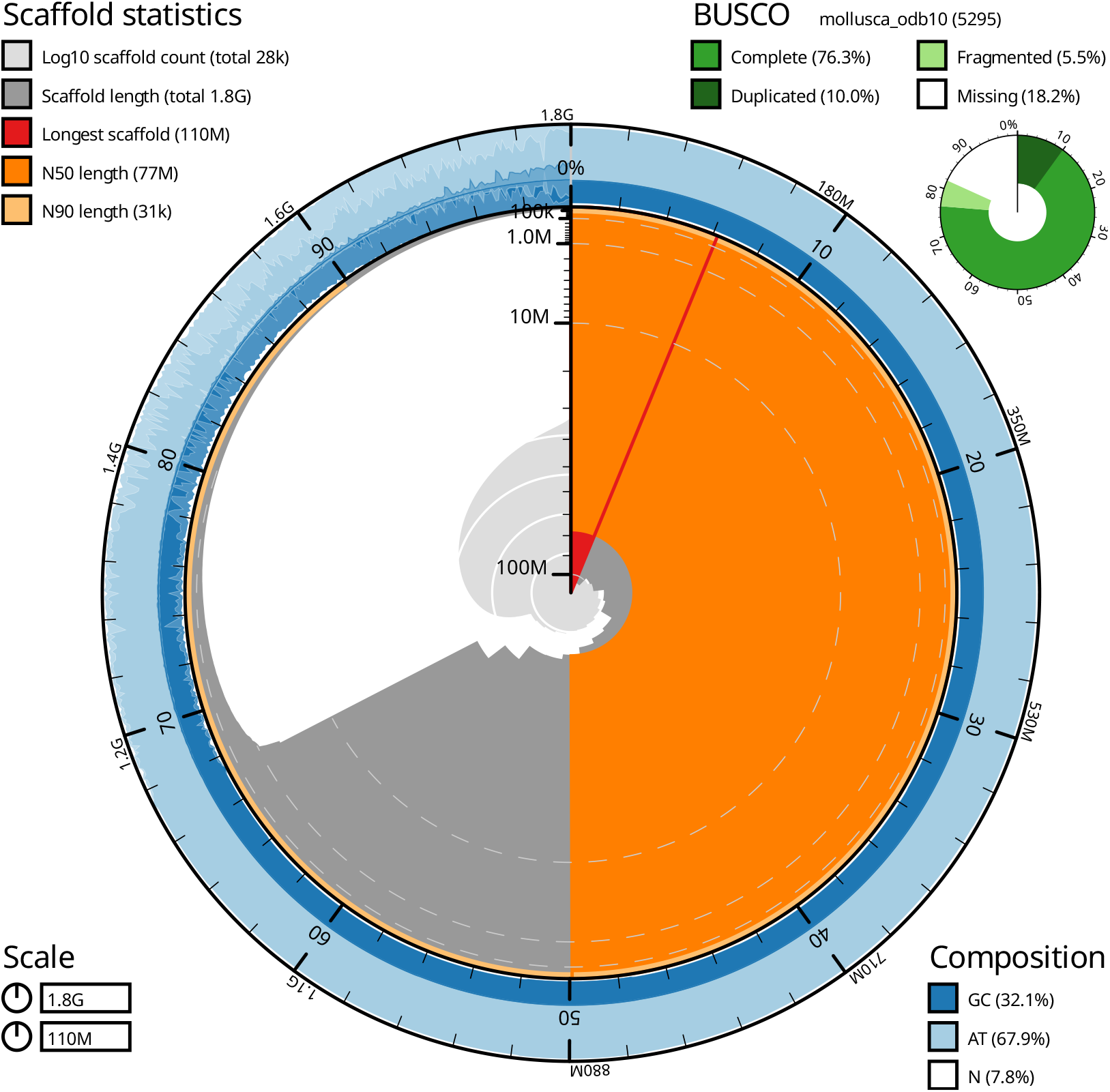
Blobtoolkit snail plot summary of assembly statistics for assembly MeduEUN_v7. The main plot is divided into 1,000 size-ordered bins around the circumference with each bin representing 0.1% of the 1,764,246,486 bp assembly. The distribution of scaffold lengths is shown in dark gray with the plot radius scaled to the longest scaffold present in the assembly (110,194,685 bp, shown in red). Orange and pale-orange arcs show the N50 and N90 scaffold lengths (77,102,752 and 30,649 bp), respectively. The pale gray spiral shows the cumulative scaffold count on a log scale with white scale lines showing successive orders of magnitude. The blue and pale-blue area around the outside of the plot shows the distribution of GC, AT and N percentages in the same bins as the inner plot. A summary of complete, fragmented, duplicated and missing BUSCO genes in the mollusca_odb10 set is shown in the top right.

**Table S2:**
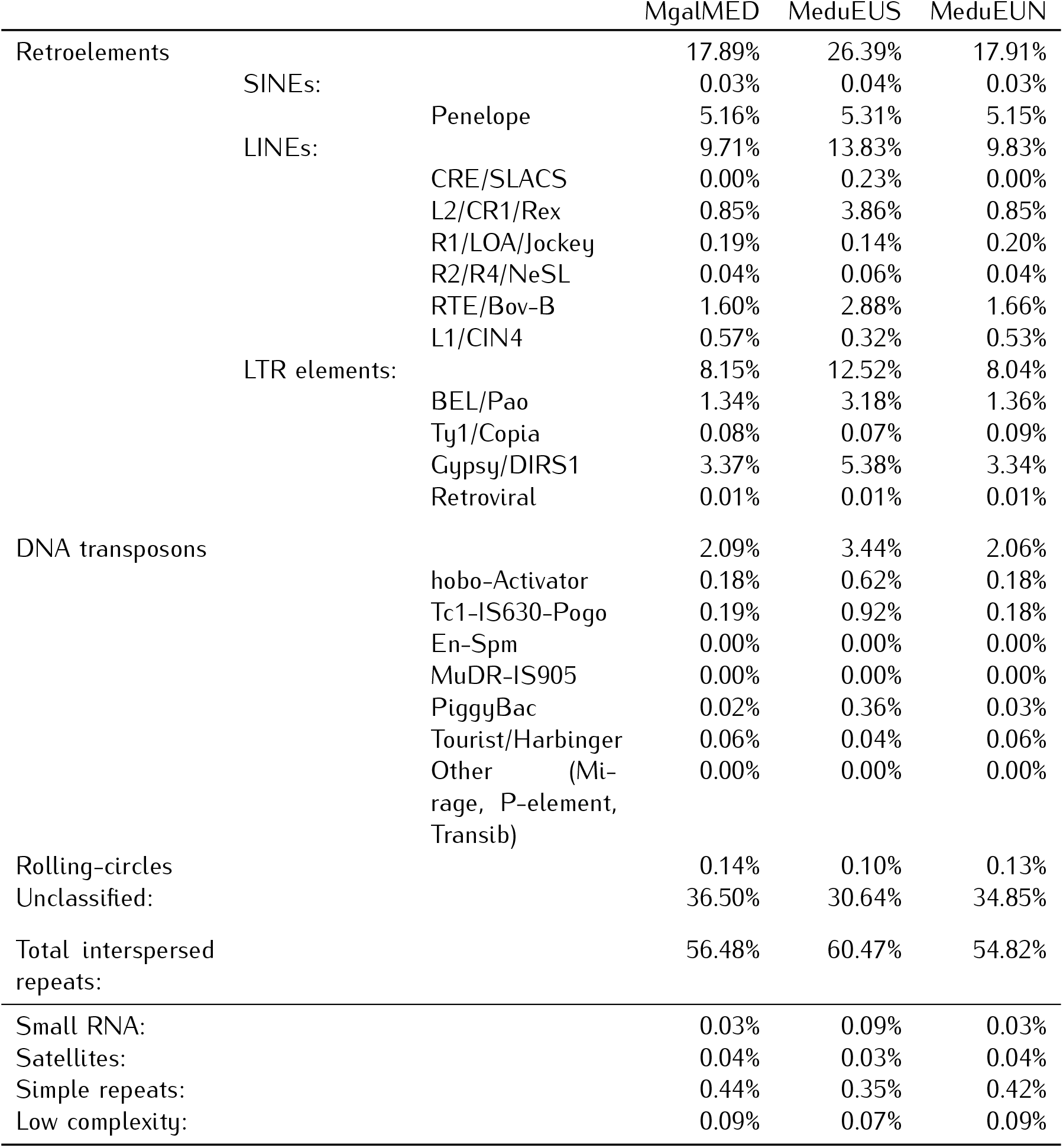
RepeatMasker results.

